# Extracellular vesicle-mediated RNA release in *Histoplasma capsulatum*

**DOI:** 10.1101/570291

**Authors:** Lysangela R. Alves, Roberta Peres da Silva, David A. Sanchez, Daniel Zamith-Miranda, Marcio L. Rodrigues, Samuel Goldenberg, Rosana Puccia, Joshua D. Nosanchuk

**Affiliations:** Instituto Carlos Chagas, Fiocruz, Cidade Industrial de Curitiba, Curitiba, Brazil; Departamento de Microbiologia, Imunologia e Parasitologia da Escola Paulista de Medicina, Universidade Federal de São Paulo - UNIFESP, São Paulo, Brazil; Departments of Medicine (Division of Infectious Diseases) and Microbiology and Immunology, Albert Einstein College of Medicine, Bronx, New York, USA; Instituto de Microbiologia da Universidade Federal do Rio de Janeiro, Brazil.

**Author notes:** Correspondence; Lysangela R. Alves, Instituto Carlos Chagas, Fiocruz, Cidade Industrial de Curitiba, Curitiba, Brazil,; Joshua D. Nosanchuk, Albert Einstein College of Medicine, 1300 Morris Park Avenue, Bronx, NY 10461, USA.

## Abstract

Eukaryotic cells, including fungi, release extracellular vesicles (EVs). These lipid bilayered compartments play essential roles in cellular communication and pathogenesis. EV composition is complex and includes proteins, glycans, pigments, and RNA. RNA classes with putative roles in pathogenesis have been described in EVs produced by fungi. Here we describe the RNA content in EVs produced by the G186AR and G217B strains of *Histoplasma capsulatum*, an important human fungal pathogen. A total of 124 mRNA were identified in both strains. In this set of RNA classes, 93 transcripts were enriched in EVs from the G217B strain, while 31 enriched in EVs produced by the G186AR strain. This result suggests that there are important strain-specific properties in the mRNA composition of fungal EVs. We also identified short fragments (25-40 long) that were strain-specific, with a greater number of them identified in EVs produced by the G217B strain. Remarkably, the most enriched processes were stress responses and translation. Half of these fragments aligned to the reverse strand of the transcript, suggesting the occurrence of miRNA-like molecules in fungal EVs. We also compared the transcriptome profiles of *H. capsulatum* with the RNA composition of EVs and no correlation was observed. Altogether, our study provided information about the RNA molecules present in *H. capsulatum* EVs, and the differences in composition between the G186AR and G217B strains. In addition, we showed that the correlation between the most expressed transcripts in the cell and their presence in the EVs, reinforcing the idea that the RNAs were directed to the EVs by a regulated mechanism.

**Importance:** Extracellular vesicles (EVs) play important roles in cellular communication and pathogenesis. The RNA molecules in EVs have been implicated in a variety of processes. In pathogenic fungi, EV-associated RNA classes have recently been described; however, only a few studies describing the RNA in fungal EVs are available. An improved knowledge on EV-associated RNA will contribute to the understanding of their role during infection. In this study, we described the RNA content in EVs produced by two isolates of *Histoplasma capsulatum*. Our results add this important pathogen to the current short list of fungal species with the ability to use EVs for the extracellular release of RNA.

## Introduction

*Histoplasma capsulatum* is major human fungal pathogen on the global stage that causes disease in both immunocompetent and immunocompromised individuals, albeit the risk for severe disease increases with compromised immunity (e.g. in patients with HIV or cancer as well as individuals receiving steroids or TNF-alpha blockers). In the United States of America, it is the most common cause of fungal pneumonia (1). *H. capsulatum* is a particular concern in certain developing regions (2), especially in Latin American countries including Brazil (3, 4), Guatemala (5), and French Guiana, where it is considered the “first cause of AIDS-related death” (6). Despite its clear importance, enormous gaps exist in our understanding of the pathogenesis of histoplasmosis, the disease caused by *H. capsulatum.* An interesting facet of *H. capsulatum’s* biology is its ability to release extracellular vesicles (EVs) (7, 8).

EVs are bilayered lipid structures released by remarkably diverse cells across all kingdoms (9). We have demonstrated that EVs are present in both ascomycetes and basidiomycetes (7, 10–14). This observation implies that mechanisms for EV production and release are truly ancient, as they appear to predate the divergence of these branches 0.5–1.0 billion years ago. Fungal EVs can carry biologically active proteins, carbohydrates, lipids, pigments and nucleic acids (15, 16), many of which are constituents of the fungal cell wall and diverse others are associated with stress response and pathogenesis.

EV-mediated transport of fungal RNA was recently shown in both commensal and opportunistic fungi. EV RNA molecules, mostly smaller than 250 nt, were identified in *Cryptococcus neoformans, Paracoccidioides brasiliensis, Candida albicans, Saccharomyces cerevisiae,* and *Malassezia simpodialis* (17, 18). Since *H. capsulatum* packages diverse compounds within EVs, we postulated that it too would use these compartments to export RNA. In this study, the EV-associated RNA components were characterized in two different isolates of *H. capsulatum*. As described in other fungi, *H. capsulatum* EVs carry both mRNAs and non-coding (nc)RNAs. In addition, proteomic data allowed the identification of 139 RNA-binding proteins in the EVs, suggesting that proteins involved in RNA metabolism might play an important role in cell communication through the EVs. Our results add this important pathogen to the list of fungal species with the ability to use EVs for the extracellular release of RNA.

## Results

### *Histoplasma capsulatum* EVs contain RNA

We characterized the RNA molecules contained in EVs isolated from culture supernatant samples of the *H. capsulatum* strains G186AR and G217B. These strains belong to distinct clades, and G217B is more virulent than G186AR in experimental models (19, 20). The most well-known difference between these two strains is that G217B lacks alpha-1,3-glucan on the yeast form cell wall (19, 20).

The reads obtained from the mRNA libraries (reads >200 nt) were aligned with each strain-specific genome available at the NCBI (G186AR ABBS02 and G217B ABBT01. For data validation, we only considered sequences with expression values of Transcripts Per Million (TPM) ≥ 100 in all biological replicates and transcripts with reads covering at least 50% of the CDS. The sRNA fraction was analyzed for the presence of different species of non-coding (nc)RNA by aligning the small RNA fraction (reads <200 nt) with the *H. capsulatum* G186AR strain. These RNA molecules were compared between the strains in order to gain insights into the role of the EV-RNA in this fungus and also to determine if there were differences in their composition between the two strains with distinct phenotypes.

### Strain-specific content of EV RNA in *H. capsulatum*

We identified a total of 124 mRNA sequences in EV samples from the two strains and carried out paired comparison between the G186AR and G217B samples. We applied the statistical negative binomial test with filters corresponding to TPM ≥ 100, log2 ≥ 2 and FDR ≤ 0.05. We observed 93 transcripts enriched in EVs derived from the G217B strain, while 31 transcripts were enriched in the G186AR strain (Supplemental Table 1). From the G217B-associated transcripts, we observed enrichment in biological processes for vesicle-mediated transport (18%), oxidation-reduction mechanisms (12%), transmembrane transport (11%) and translation (8%) (Figure 1). For the G186AR strain, the mRNA sequences were only enriched in general cellular and metabolic processes (59%). These results suggest that there are important differences in the mRNA composition of EVs derived from these two strains of *H. capsulatum*.

**Figure 1:**
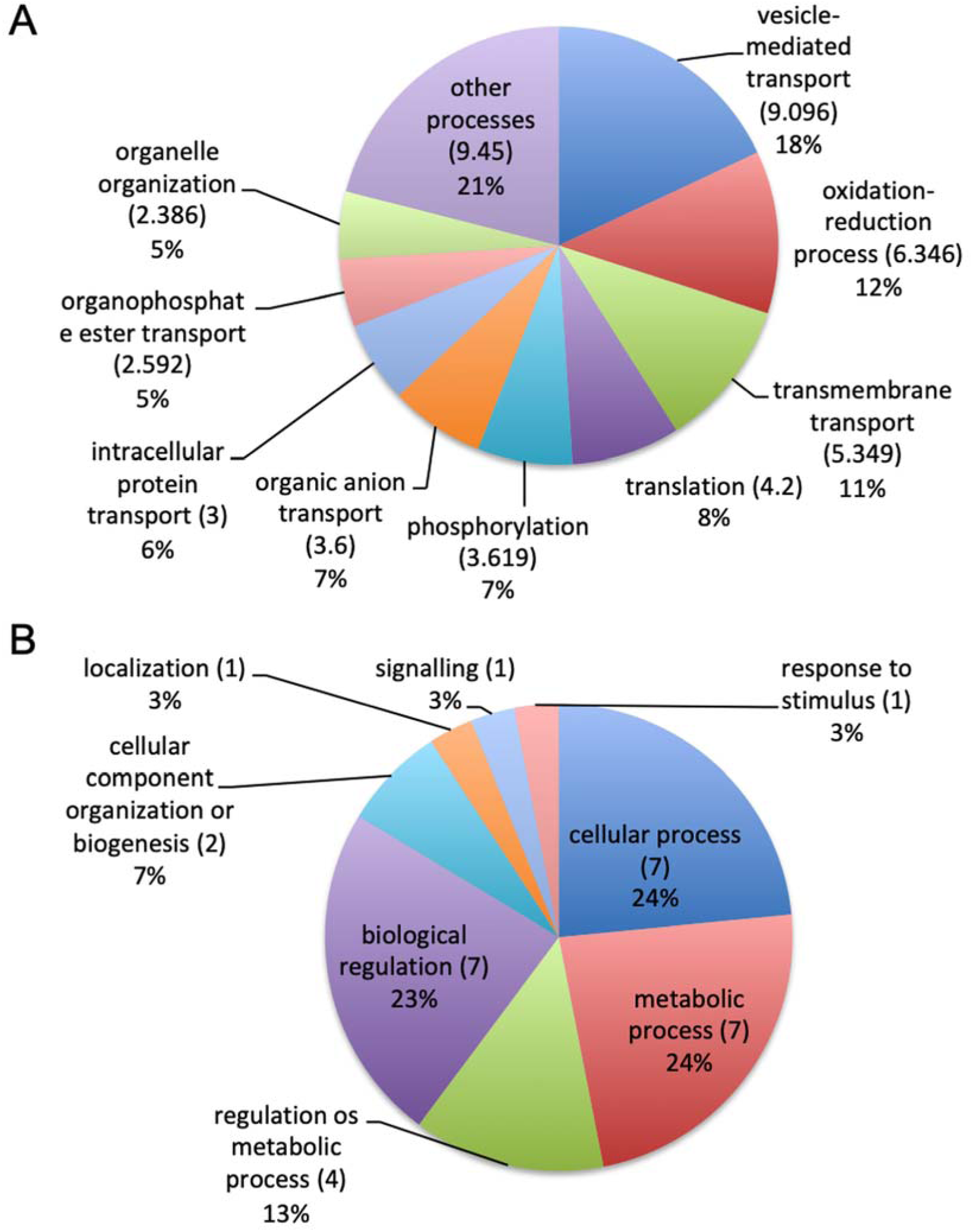
Gene ontology analysis. Pie chart representing the gene ontology of mRNA sequences enriched in EVs isolated from A) *H. capsulatum G*217B, n = 93. B) *H. capsulatum G*186AR, n=31.

### *H. capsulatum* EVs contain mRNA fragments and miRNA-like molecules

In addition to the identification of full-length transcripts in EVs, we also detected short reads of 25-40 nt in average that aligned consistently in the CDS, but at specific positions of the mRNAs (3’, 5’or middle); about 50% of these short fragments aligned to the reverse strand. A total of 172 (G217B), and 80 (G186AR) sequences of this type (Table 1). A total of 172 fragments were represented in the G217B sample compared to only 80 found in the G186AR EVs (Table 1). About 47% of the reference mRNA translate proteins of unknown biological processes. Those associated with DNA metabolism/biogenesis were the second most abundant for both EV samples (22 for G217B versus 16 for G186AR), followed by transport for G217B, and protein modification for both strain EVs. Other processes related to short RNAs identified in both strain EVs were oxidation-reduction, signaling, and carbohydrate and lipid metabolism (Table 1). RNA fragments associated with translation were highly enriched in G217B (11) but not in G186AR (2) EVs, while those related to response to stress were found exclusively in the G217B sample. The corresponding proteins are stress response protein whi2, the DNA repair protein rad5 and a thermotolerance protein (Table 1). Analysis of translation-related sequences allowed identification of mRNA fragments associated to distinct steps of the translation process, such as ribosome biogenesis and processing. Other metabolic pathways identified in both strains were protein modification, carbohydrate, and lipid metabolism, signaling, oxidation-reduction and transmembrane-transport, among others (Table 1).

**Table 1:**
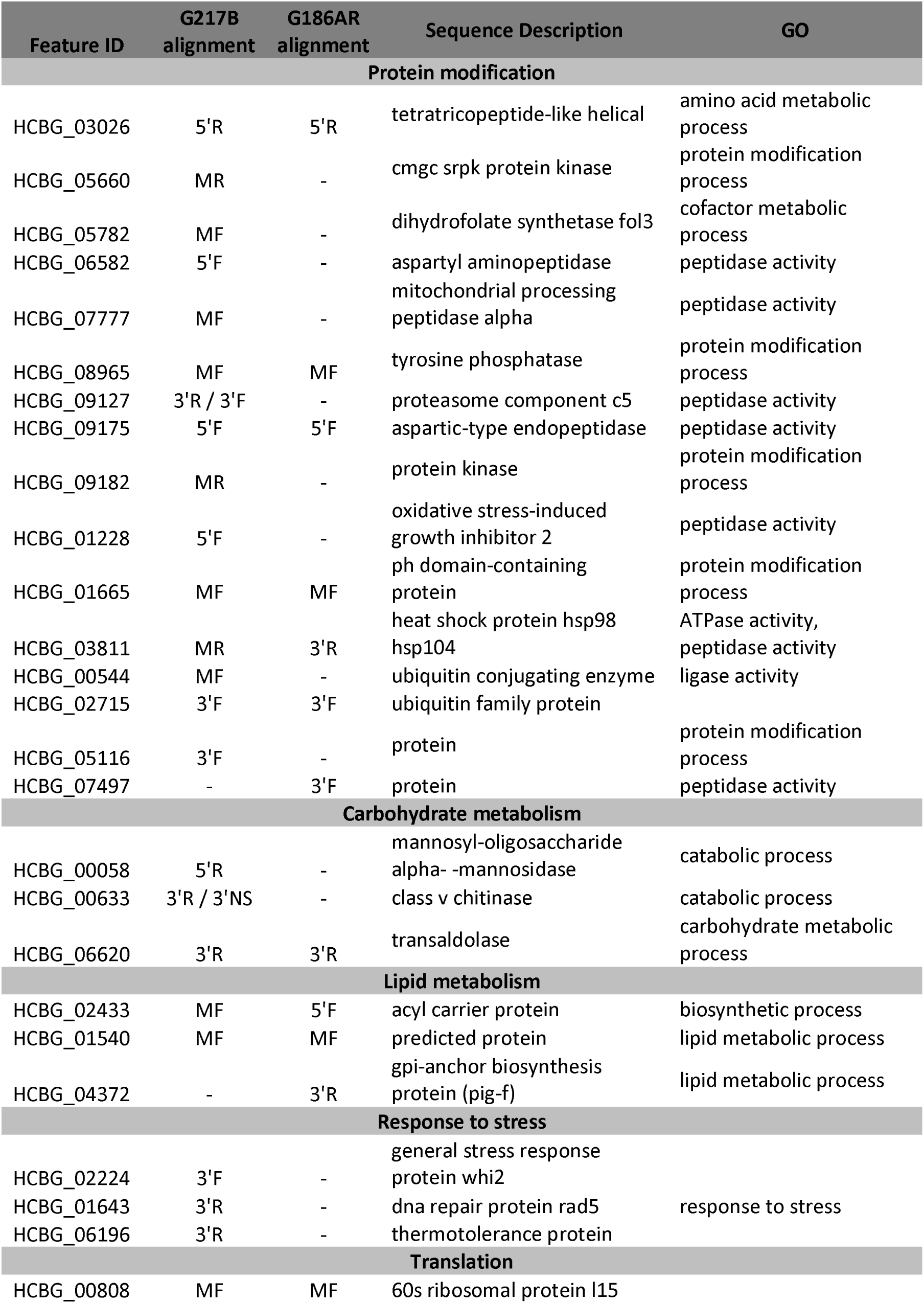
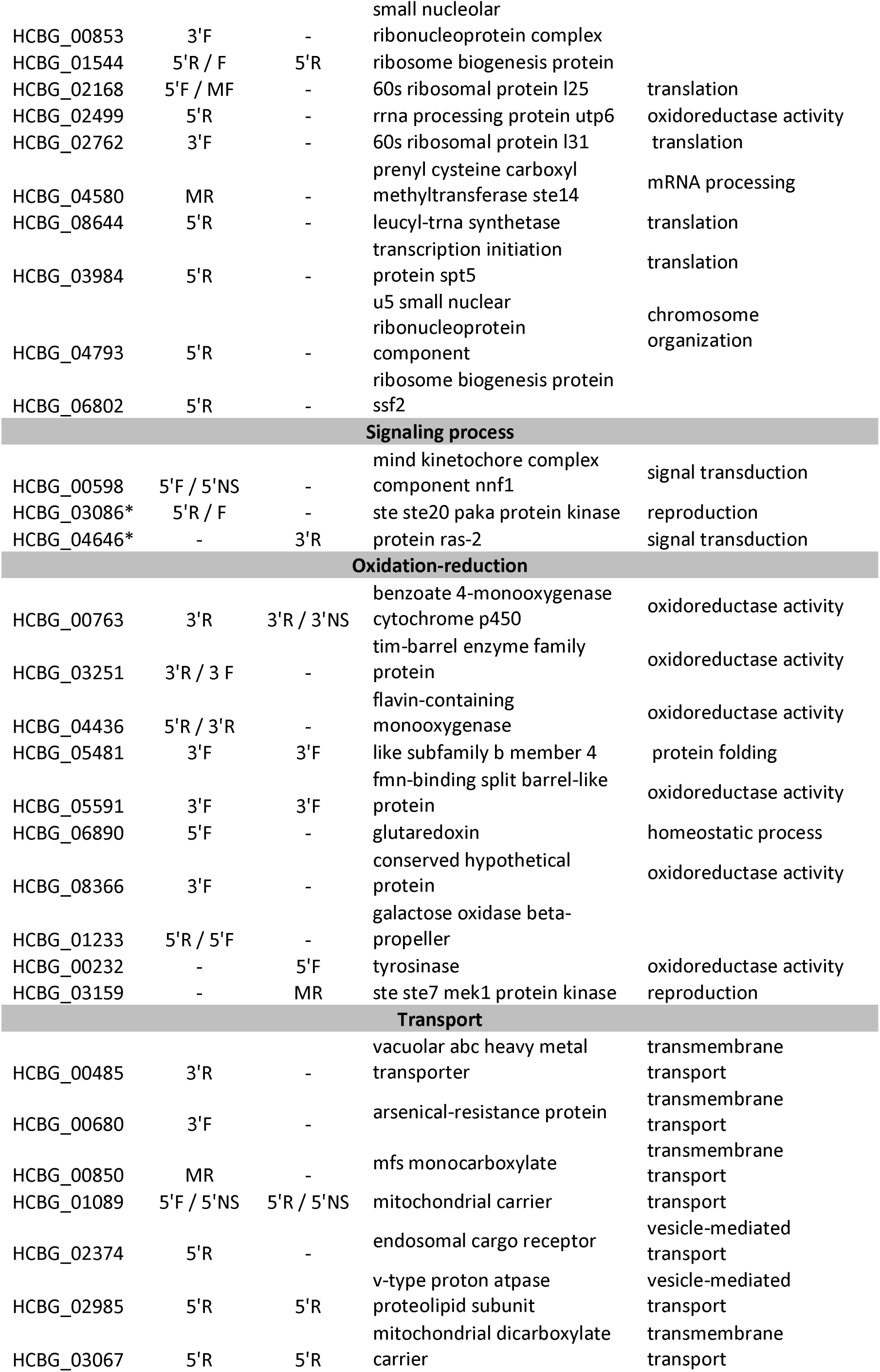
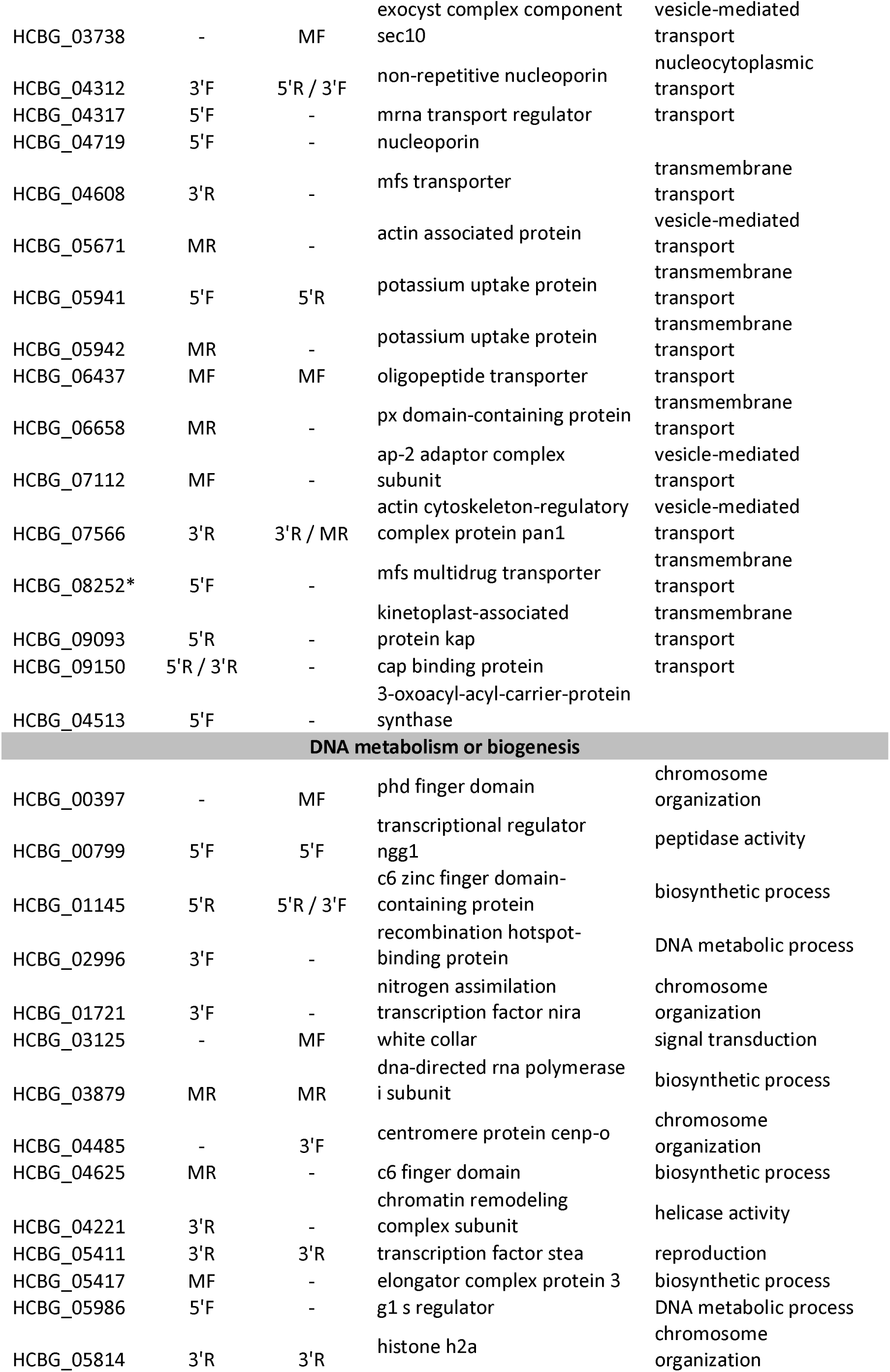
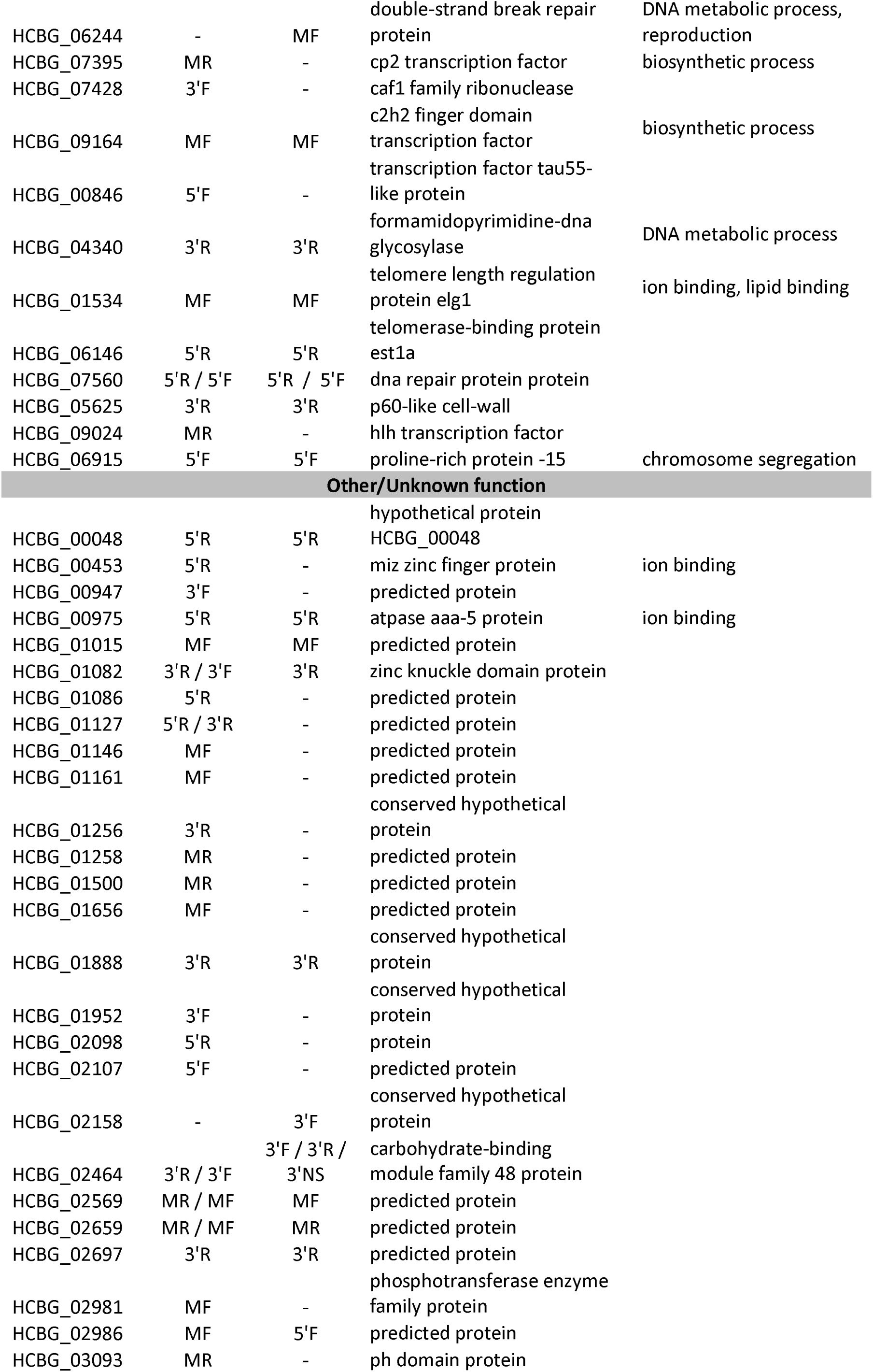
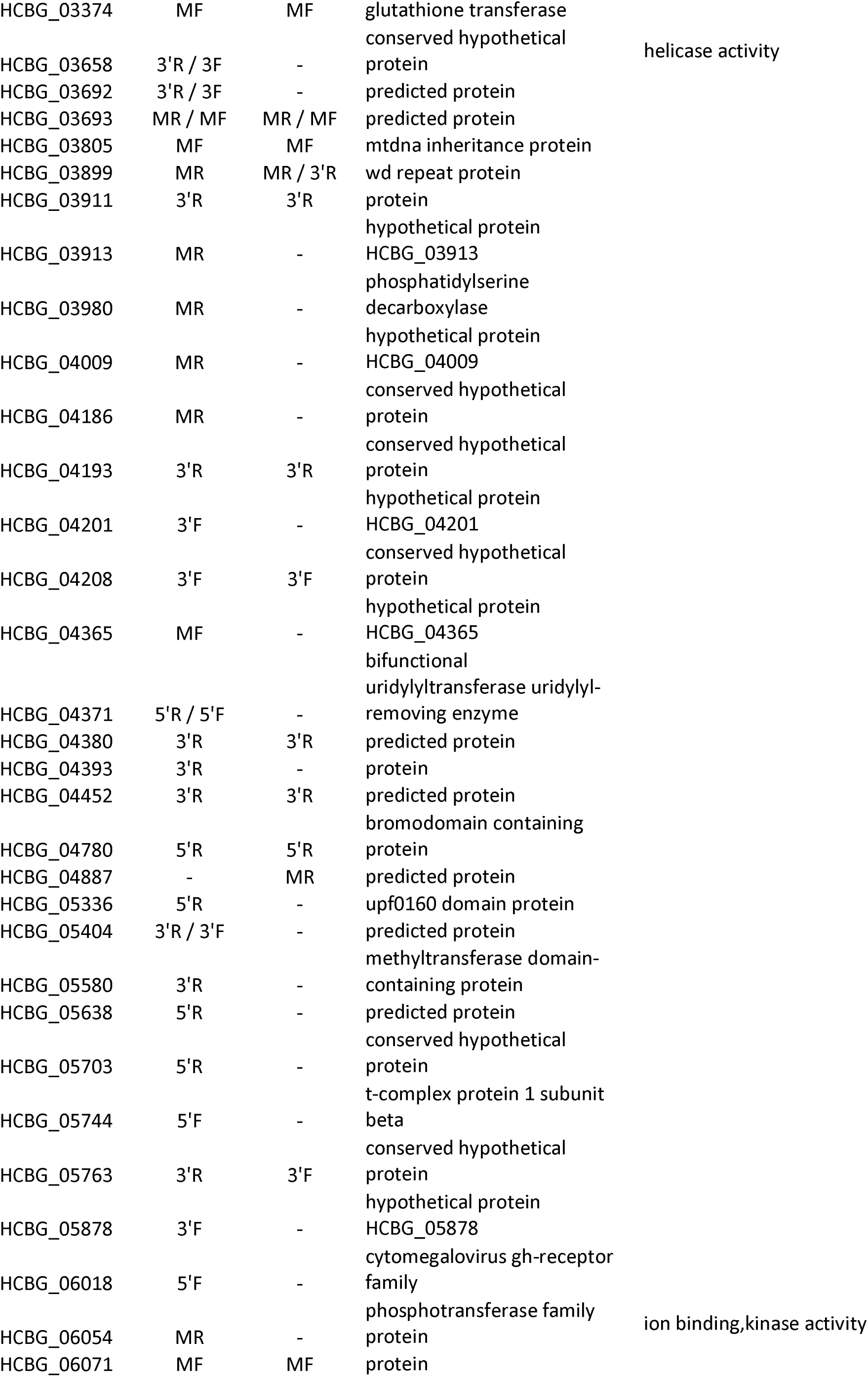
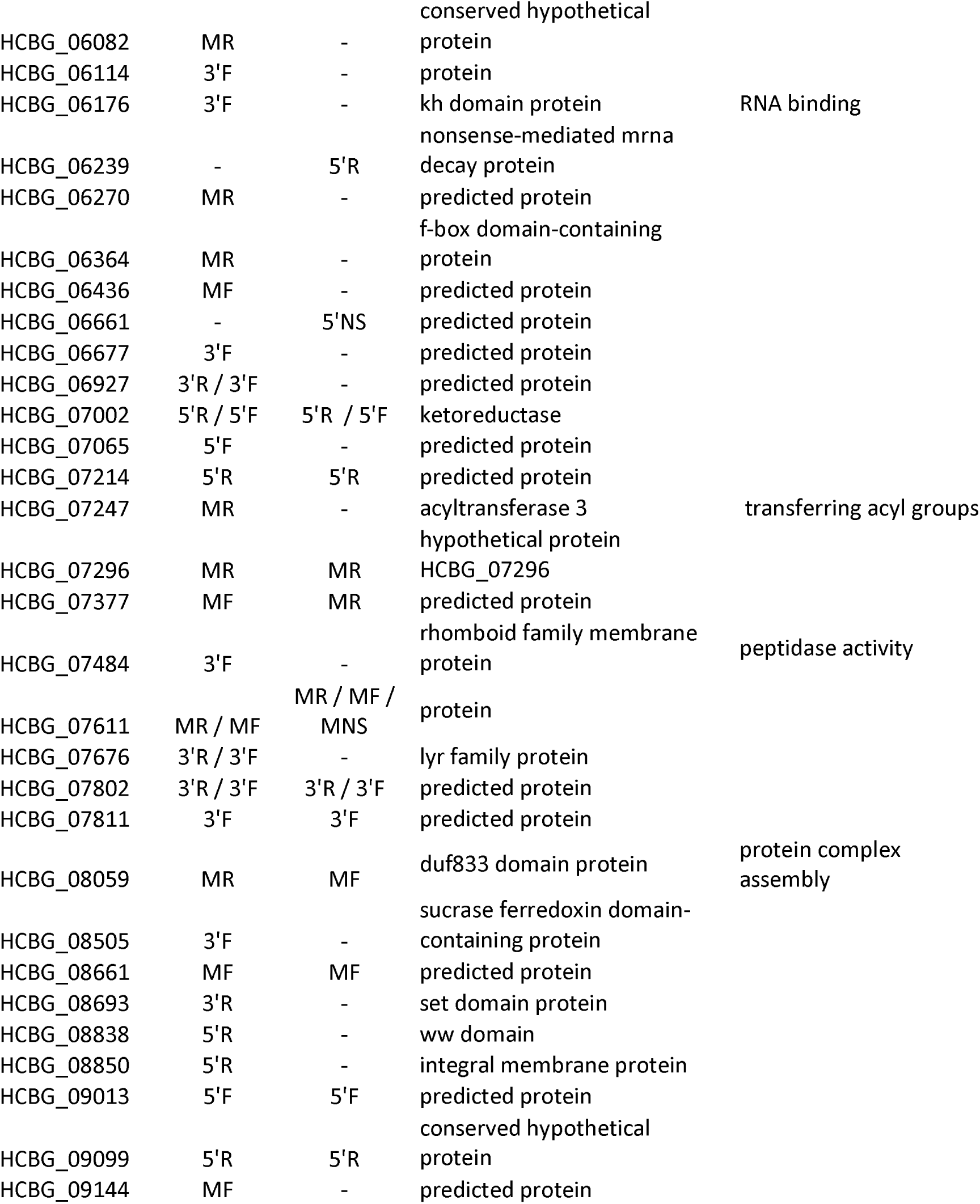
Fragments of mRNAs identified in the EVs isolated from the G217B and G186AR strains. For some transcripts, there was an alignment in specific positions of the mRNA, not covering the entire sequence. 5’, 3’ or M (middle of the mRNA); F or R orientation.

To gain further insight into the role of these mRNA-fragments, to determine if they could be derived from a miRNA-like pathway and to assess if they could play a biological role in the recipient cell, we searched for RNA secondary structures, since they are fundamental for gene expression regulation (21). A wide study of RNA structures in distinct cells revealed regulatory effects of the RNA structure throughout mRNA life cycle such as polyadenylation, splicing, translation, and turnover (22, 23). A total of 54 RNAs with putative structures were generated by a probability distribution, using a free energy (ΔG) less than or equal to – 7.0 (Supplemental table 2). On the basis of this parameter, we identified transcripts for U3 small nucleolar RNA-associated protein, L-isoaspartate O-methyltransferase, serine/threonine-protein kinase, proteasome component C5, pre-rRNA processing protein Utp22, C-x8-C-x5-C-x3-H zinc finger protein, fungal specific transcription factor domain-containing protein and DNA damage-responsive transcriptional repressor RPH1 were identified (Figure 2 and Supplemental table 2).

**Figure 2:**
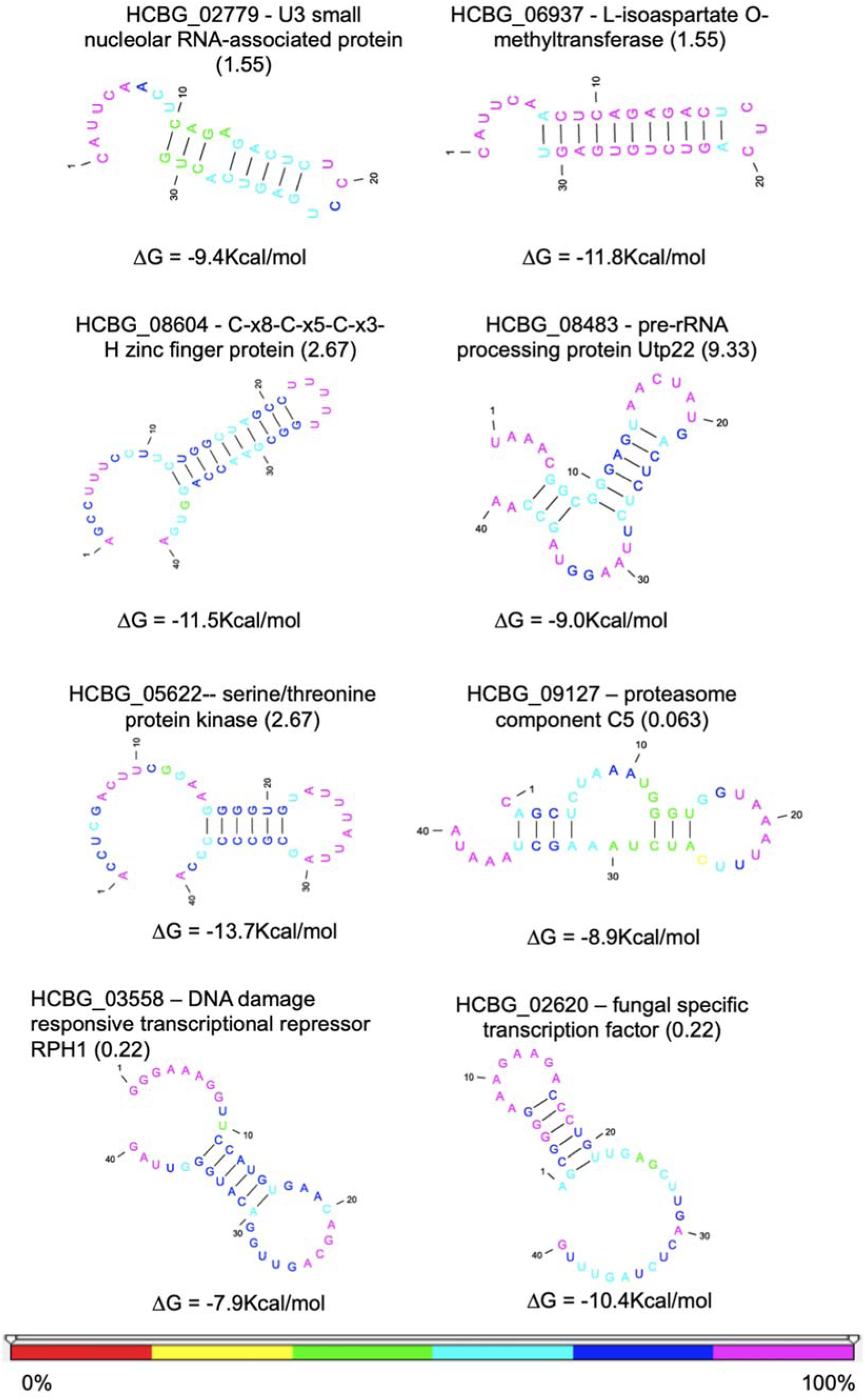
RNA secondary structure. We used the ppFold software to predict the secondary structure from the putative miRNAs-like extracted from the obtained reads. The numbers in parenthesis represent the alignment E-value. The nucleotide colors represent the reliability percentage for each position of the RNA molecule (bottom figure). The stability value of each structure is given in kcal/mol.

### Comparison of EV ncRNA classes in *H. capsulatum* EVs

We used the ncRNA database from *H. capsulatum* to identify the classes of ncRNA present in EVs RNA. The data analysis revealed 73 different sequences of ncRNA in *H. capsulatum* EVs from the G186AR strain and 38 from the G217B isolate. Thirty three molecular species were common to both strains and 40 were exclusively identified in the G186AR strain and the most abundant class of ncRNA found in *H. capsulatum* EVs was tRNAs (Table 2).

**Table 2:**
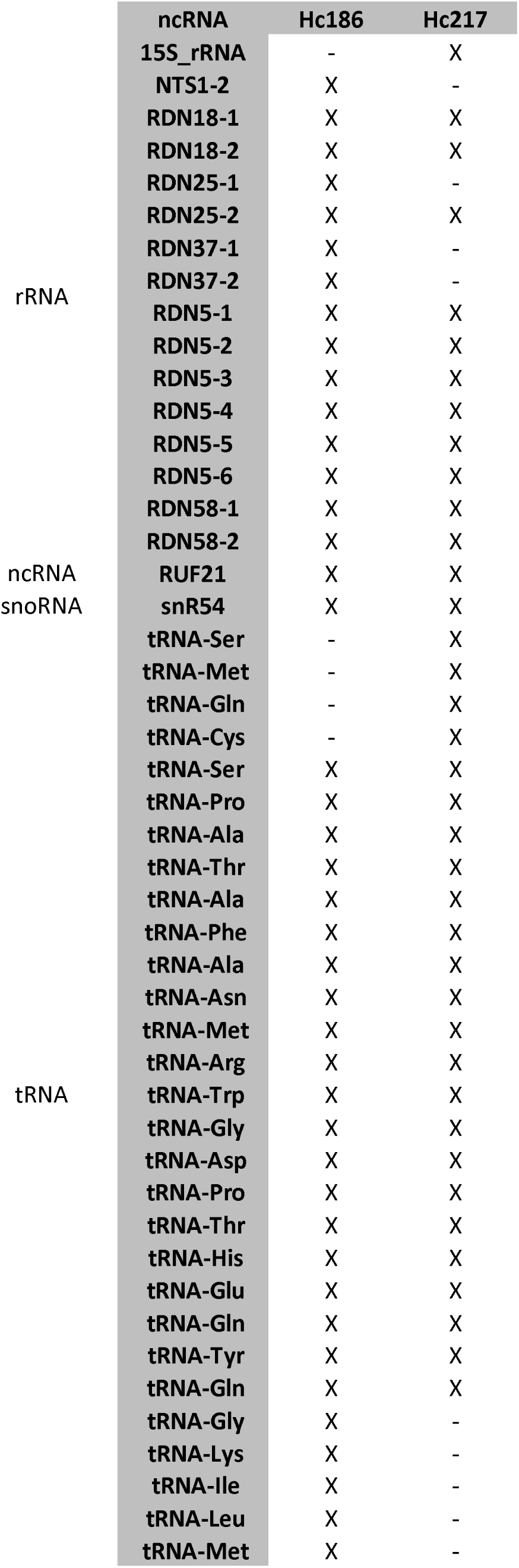
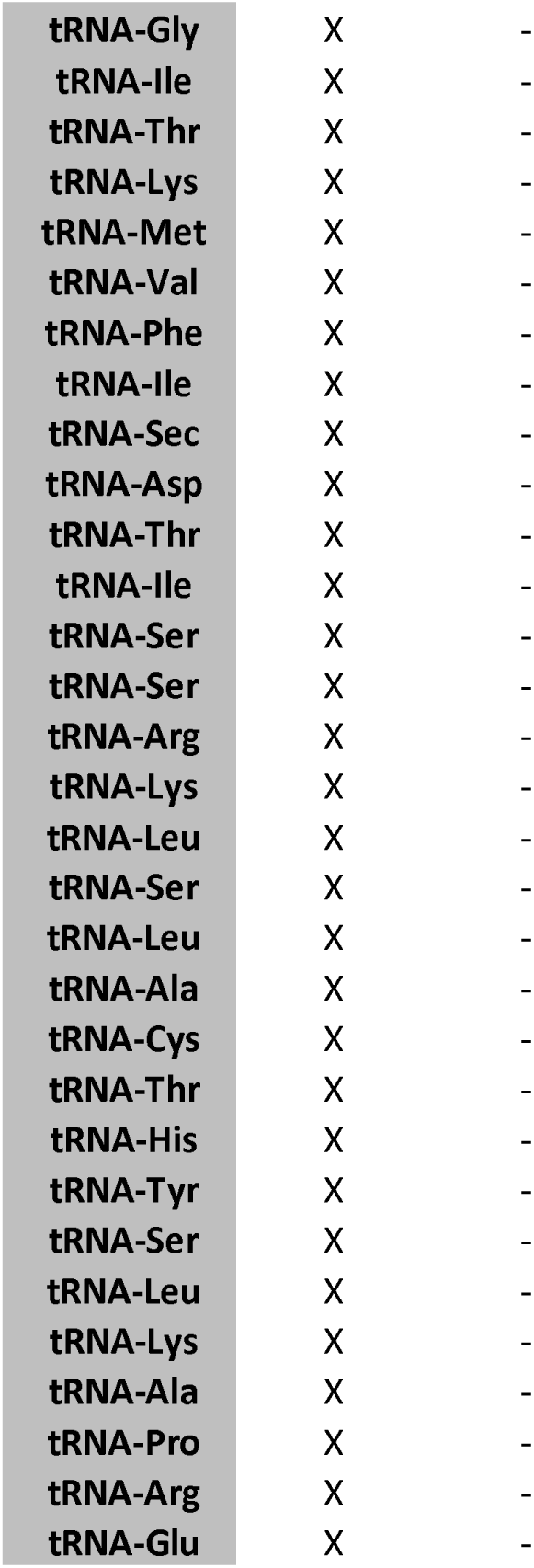
Classes of ncRNA sequences identified in EV preparations from *H. capsulatum* strains G186AR and G217B.

### Analysis of proteins putatively associated to RNA metabolism in the EVs

As a rule, cellular RNAs are covered with proteins and exist as ribonucleoprotein complexes. The proteins associated to RNAs are named RNA-binding proteins (RBPs). These proteins participate on several biological processes, from transcription to RNA decay (24). In this context, we investigated the presence of RBPs in the *H. capsulatum* EVs. We analyzed the proteomic EV data available for the G217B strain (25) and we identified 139 proteins related to RNA metabolism (8) (Table 3 and Supplemental table 3). We found many RBPs, such as PolyA binding protein (PABP), Nrd1, Prp24, and Snd1; splicing factors, exosome complex components and ribosomal proteins (Table 3 and Supplemental table 3) were identified. In addition, we also found the quelling deficient protein 2 (QDE2), an argonaute protein important in the RNA machinery in fungi. As we identified the QDE2 in EVs, we searched for the components of the RNAi machinery in *H. capsulatum*, and compared them with the proteins from *Neurospora crassa* and *Schizosaccharomyces pombe*, which are fungal species where the RNAi machinery has been most well described (26, 27). *H. capsulatum* EVs contained one argonaute protein (QDE2), two dicer-like proteins, the QIP (quelling interaction protein) and the RNA-dependent RNA polymerase (QDE1) (Table 4).

**Table 3:**
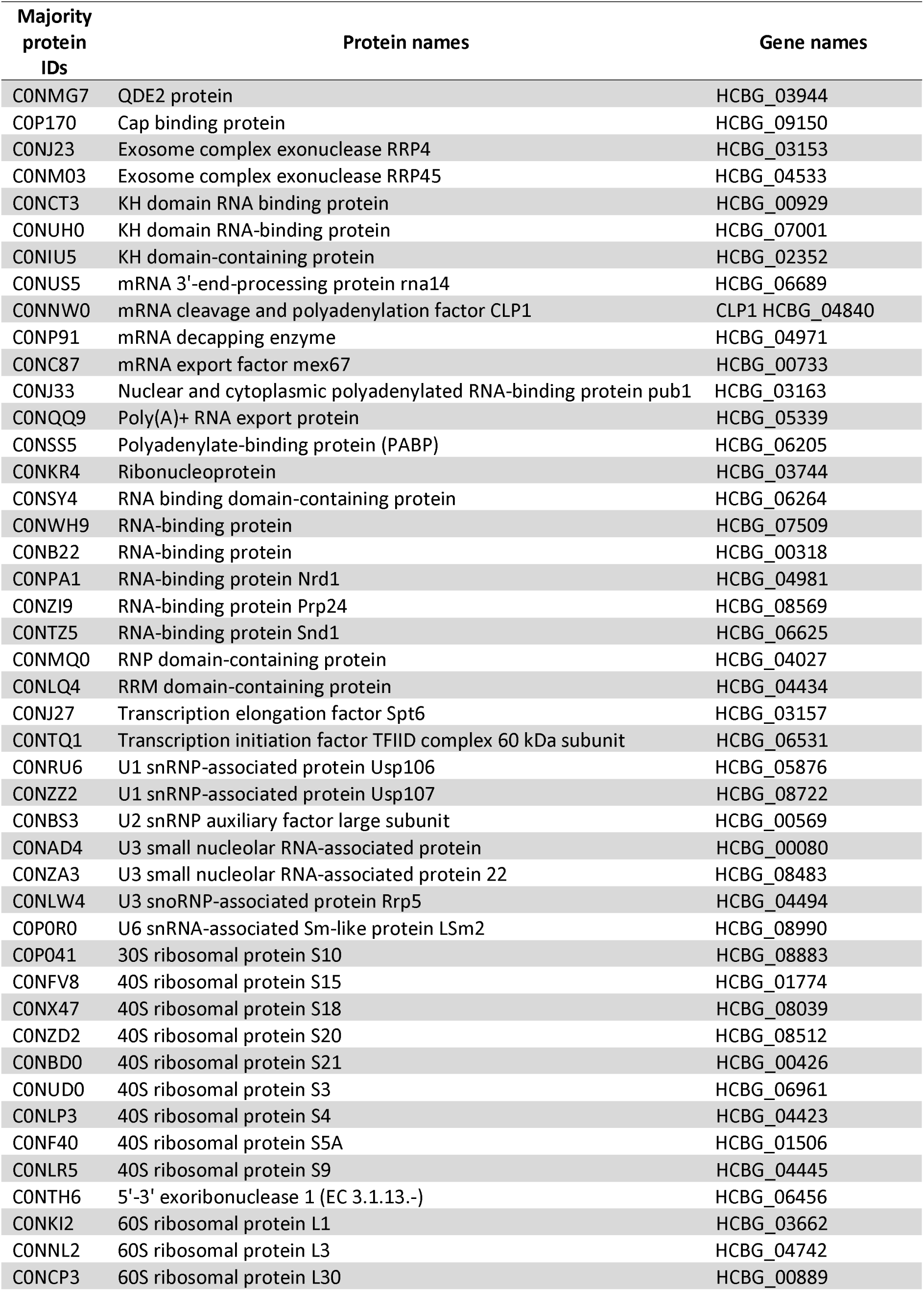
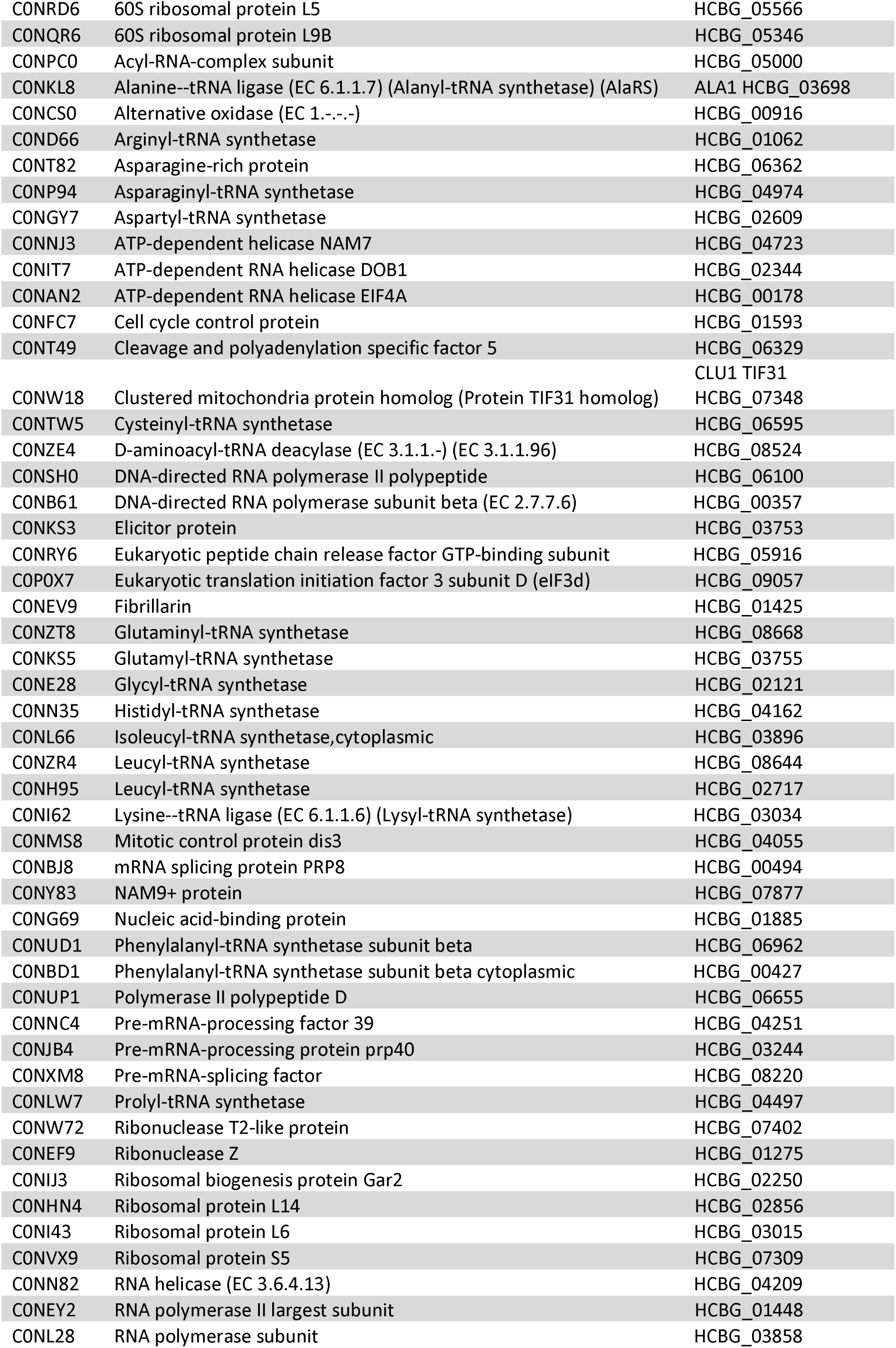
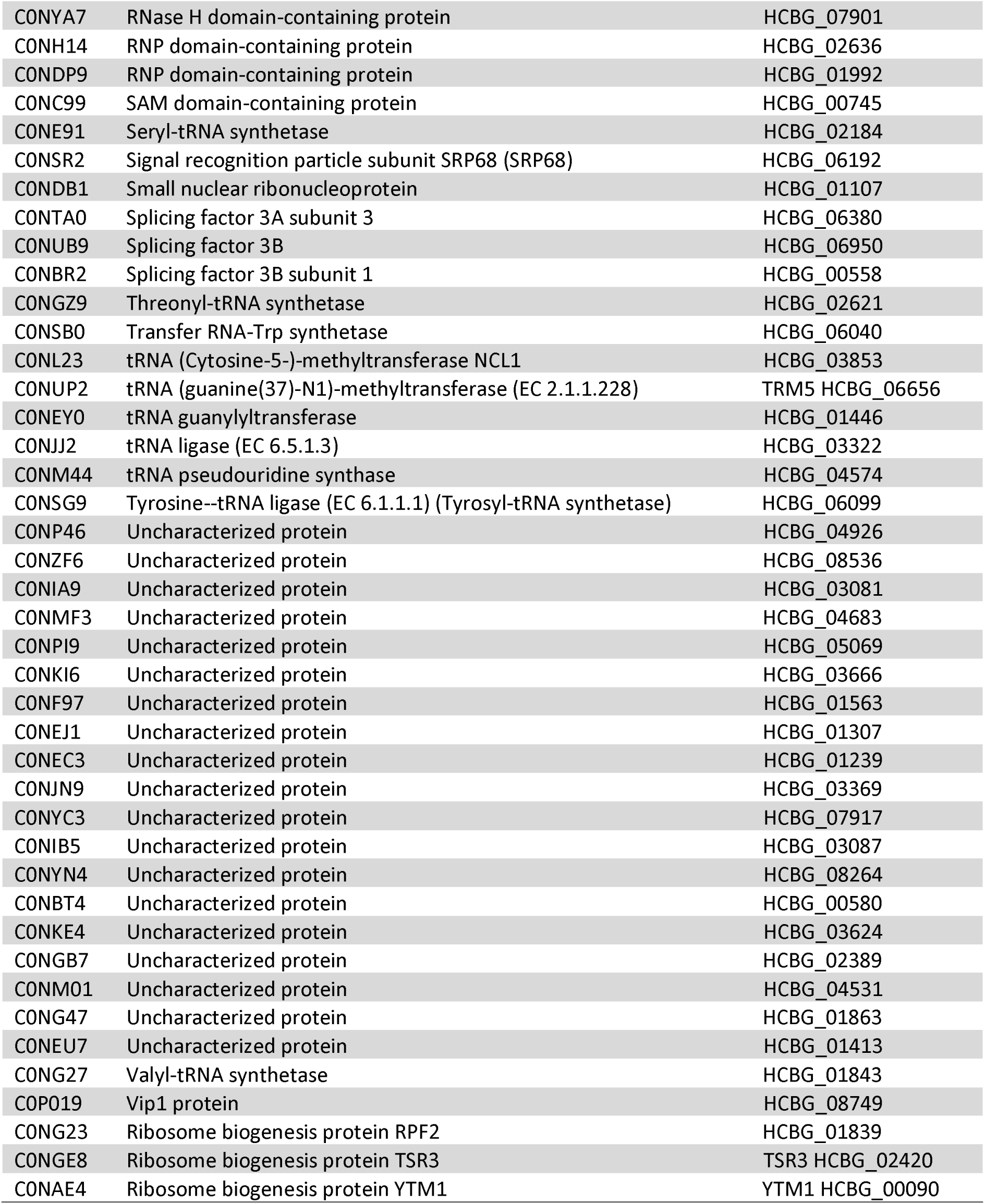
Proteins related to RNA metabolism identified in EV preparations from *H. capsulatum* strain G217B.

**Table 4:**
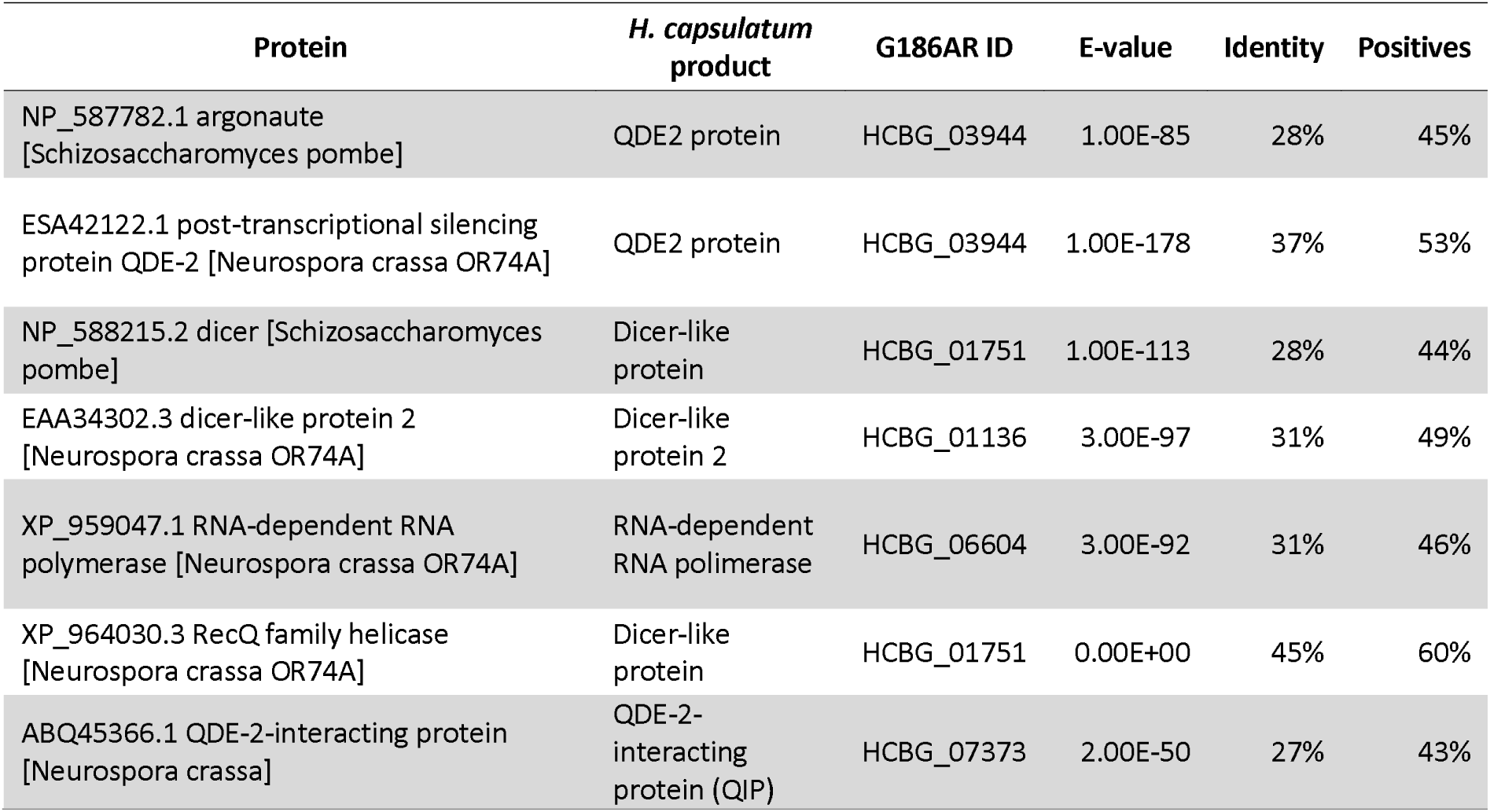
Proteins associated to the RNAi machinery in *H. capsulatum* G186AR EVs compared to *S. pombe* and *N. crassa*.

### Comparison of cellular RNA vs. EV RNA shows a distinct enrichment of molecules in the vesicles

We next assessed the composition of cellular RNA from *H. capsulatum* yeast cells (28) and compared this information to that obtained from EV-associated RNA composition under the same conditions. There was no correlation between the transcripts with highest expression levels and their presence in the EVs (Supplemental table 4). Examples of highly expressed cellular transcripts included histones 4, 2B, and 2A, allergen Aspf4, chaperones, and translation factors, among others (Supplemental table 4). In contrast, zinc knuckle domain-containing protein, vacuolar ATP synthase subunit C, G1/S regulator, thermotolerance protein, histone variant H2A.Z and proteasome component C5 had an enrichment value greater than 7,000 in the EVs, while they showed low expression values in the cell (Supplemental table 4). The differences in composition between cells and EVs were also evaluated by grouping the transcripts into biological processes (Figure 3). For the yeast cells, the main pathways were associated with transport, translation and general metabolic processes (Figure 3). For the EVs, the enriched pathways were transmembrane transport, protein phosphorylation and transcription regulation (Figure 3). This result demonstrates the low levels of correlation between the most expressed cellular mRNAs and EV cargo, evidencing there might be a mechanism directing the RNA molecules to the EVs.

**Figure 3:**
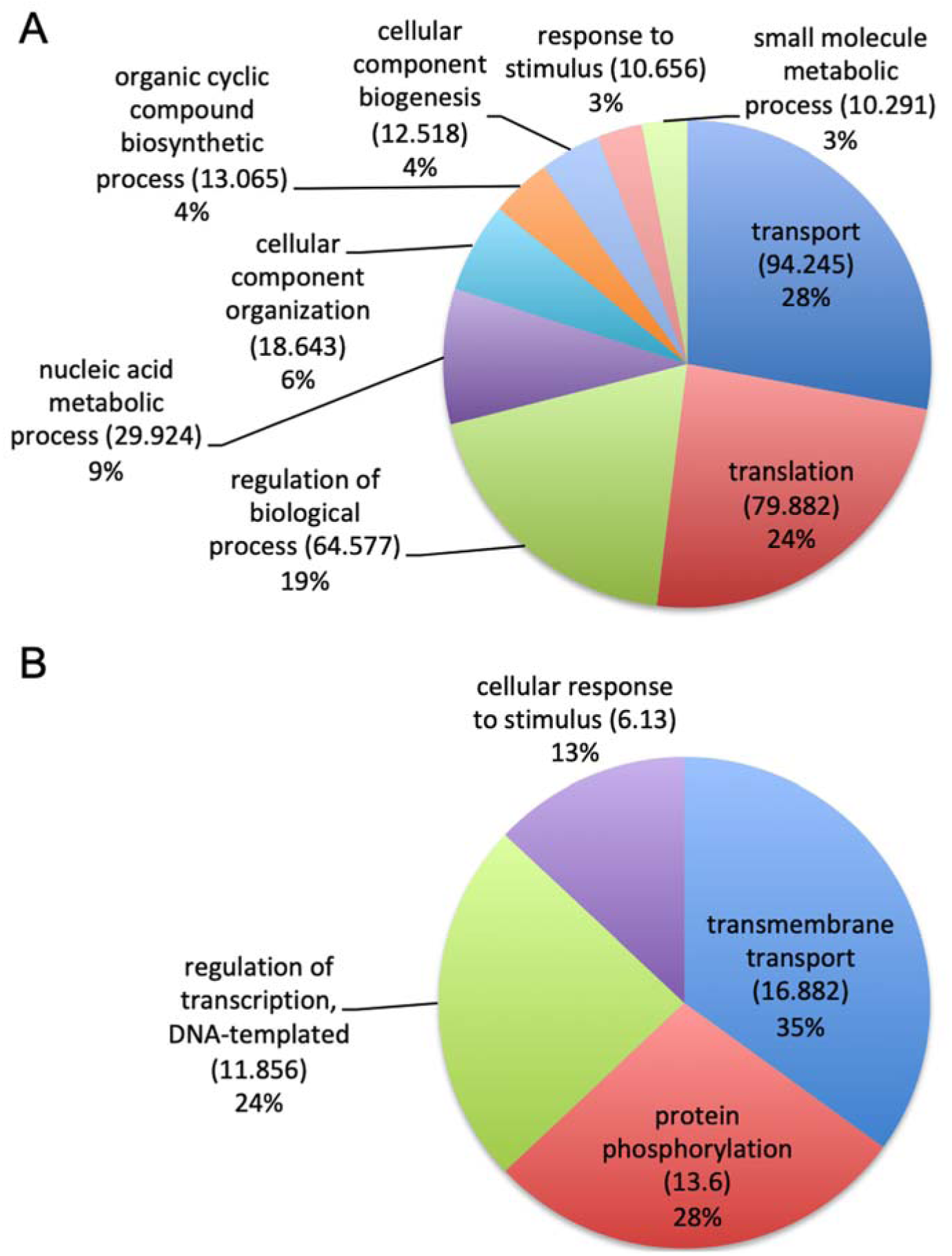
Gene ontology analysis. Pie chart representing the gene ontology of mRNA sequences enriched in A) *H. capsulatum* cells and B) in EVs isolated from *H. capsulatum*.

## Discussion

As previously described (17, 18), RNA molecules associated to fungal EVs are remarkably diverse. For instance, mRNAs, tRNA fragments, snoRNAs, snRNAs, and miRNA-like molecules were characterized in EVs from *C. albicans, C. neoformans, P. brasiliensis* and *S. cerevisiae* (17). In *H. capsulatum* EVs we observed a similar distribution of RNA molecules. The comparison between G186AR and G217B EVs revealed important differences in the variety of mRNAs identified. When the mRNA composition was compared to what was described for other fungi, important similarities were observed. For example, the most abundant biological process identified in G217B EVs was vesicle-mediated transport, which was also the most abundant process in *C. albicans* EVs (17). Molecules required for ribosome biogenesis, which were observed in G217B EVs, belonged to the most enriched process in *S. cerevisiae* EVs (17). However, when the ncRNA molecules were compared, different profiles were observed. Most of the ncRNA in *H. capsulatum* strains derived from tRNAs; a similar profile was obtained with *C. albicans* (17). In addition, in *H. capsulatum*, almost no snoRNAs were identified, but this class of ncRNAs was one of the most abundant in the EVs of other fungi (17). Differences in EVs composition have been observed in *C. neoformans;* EV-associated RNA produced by mutant cells with defective unconventional secretion differed considerably from similar samples produced by wild-type cells (29).

In our study, we identified short reads that aligned specifically to exons; however, these sequences did not correspond to complete mRNAs in the EVs. They rather corresponded to 25 nt long fragments that were enriched in specific exons of the transcript. These fragments of mRNAs were previously described in human cells (30) where most of the transcripts identified in the EVs corresponded to a fraction of the mRNA with an enrichment of the 3′-end of the transcript (30). This human study led to the hypothesis that the mRNA fragments had a role in gene expression regulation in the recipient cells as the secreted mRNA could act as competitors to regulate stability, localization and translation of mRNAs in target cells (30). In *Mucor circinelloides* cells, the RNA silencing pathway (sRNA) resulted in the production of both sense and antisense small RNAs (31–33). Sequencing analysis of the small RNA content of this fungus showed the existence of exonic small interfering RNAs (ex-siRNA) as a new type of sRNA. They were produced from exons of the same genes that are later regulated through the repression of the corresponding mRNA (34). This result agrees with our observation of short reads in the exonic regions of the transcripts. We therefore hypothesize that; similar to what was described for *M. circinelloides* cells, and the *H. capsulatum* EV fragments can regulate expression of their own mRNAs. Of note, we also found a highly represented population of putative exonic-siRNA in *Paracoccidioides* strains (Peres da Silva et al., submitted).

As *H. capsulatum* EVs contain different RNA molecules, it is reasonable to hypothesize that proteins that regulate RNA metabolism are also present in the EVs, probably associated to RNA. If validated, this hypothesis could indicate how a specific subset of RNAs are directed to the vesicles and exported. RNA binding proteins (RBPs) participate in several biological processes, from RNA transcription to decay (24). We detected a number of RNA binding proteins in *H. capsulatum* EVs (25). In other systems, these proteins were also identified in association with EVs. For example, in EVs produced by human epithelial cells, 30 RBPs were identified (35), including heterogeneous nuclear ribonucleoproteins (hnRNPs). These proteins are responsible for directing pre-mRNAs in the maturation processes that culminate with transcriptional regulation, alternative splicing, transport, and localization (35). In addition, RBPs in EVs were identified in distinct models as hepatocytes, human embryonic kidney (HEK) cells, and mouse myoblast cells (35–37). Interestingly, one of the RBPs identified in EVs was SND1 (Staphylococcal nuclease domain-containing protein 1), which is a main component of RISC complex (RNA-induced silencing complex) that plays an important role in miRNA function (37).

Another example of a protein identified in the EVs of *H. capsulatum* and distinct organisms is an endonuclease of the Ago2 family. An infection model with *Plasmodium falciparum* demonstrated that infected red blood cells released EVs containing functional miRNA-argonaute 2 complexes (38). Moreover, endothelial cells internalized the *P. falciparum* EVs, and the miRNA-argonaute 2 complex were transferred to the cells and acted regulating the gene expression and in the barrier properties of the recipient cells (38). The argonaute protein in *H. capsulatum* named QDE2 was identified enriched in the EVs of the G217B strain.

Small silencing RNAs include a variety of molecules, such as microRNAs (miRNAs) and various small interfering RNAs (siRNAs), such as exo-siRNAs, endo-siRNAs, and pi-RNAs (39). Previous studies of small RNAs in fungi have identified the RNAi machinery in the fission yeast *Schizosaccharomyces pombe*, in the budding yeast *S. castellii, C. albicans*, and in filamentous fungi (26, 27, 40). One of the best-characterized models is the filamentous fungus *N. crassa* (27, 41–45). The RNAi machinery in this organism is a defense against transposons (46). A similar process has been described in *C. neoformans*, where RNAi is involved in the regulation of transposon activity and genome integrity during vegetative growth (47). In *N. crassa,* the *QDE2* gene encodes an Argonaute protein that is homologous to the rde-1 gene in *C. elegans*, a protein required for dsRNA-induced silencing (27). The characterization of RNAs associated to QDE2 in *N. crassa* led to the identification of miRNA-like RNAs (milRNAs) in this organism (48). The identification of QDE2 in *H. capsulatum* EVs in association to the small RNAs indicate that the complex QDE2-milRNA might be directed to the EVs and possibly delivered to recipient cells, with the potential to interfere with gene expression regulation and / or cell-cell communication.

Fungal EVs have been implicated in a number of communication processes, including transfer of virulence (49) and antifungal resistance (50). In *C. gattii*, pathogen-to-pathogen communication via EVs reverted an avirulent phenotype through mechanisms that required vesicular RNA (49). The sequences required for this process, however, remained unknown. This is an efficient illustration of the potential derived from the characterization of EV-associated RNA in fungi. In this context, our study provides information in the *H. capsulatum* model that will allow the design of pathogenic experimental models aiming at characterizing the role of extracellular RNAs in fungal pathogenesis.

## Material and Methods

### Fungal strains and growth conditions

The *H. capsulatum* strains were stored long term at −80°C. Aliquots were inoculated into Ham’s F-12 media (Gibco, Cat# 21700-075) supplemented with glucose (18.2 g/L), L-cysteine (8.4 mg/L), HEPES (6 g/L) and glutamic acid (1 g/L) and cultivated with constant shaking at 150 rpm at 37°C. Viability assessments were performed using Janus green 0.02%, and all aliquots used had >99% of live yeast cells. EVs were then isolated from fungal culture supernatants as previously described (51).

### sRNA isolation

Small RNA enriched fractions were isolated with the miRNeasy mini kit (Qiagen) and were then treated with the RNeasy MinElute Cleanup Kit (Qiagen), according to the manufacturer’s protocol, to obtain small RNA-enriched fractions. The sRNA profile was assessed in an Agilent 2100 Bioanalyzer (Agilent Technologies).

### RNA sequencing

One hundred ng of purified sRNA were used for RNA-seq analysis from two independent biological replicates. The RNA-seq was performed in a SOLiD 3 plus platform using the RNA-Seq kit (Life Science) according to the manufacturer’s recommendations.

### *In silico* data analysis

The sequencing data were analyzed using the version 10.1 of CLC Genomics Workbench©. The reads were trimmed on the basis of quality, with a threshold Phred score of 25. The reference genomes used for mapping were obtained from the NCBI database (*H. capsulatum* G186AR strain - ABBS02, and G217B strain – ABBT01). The alignment was performed as follows: additional 100-base upstream and downstream sequences; 10 minimum number of reads; 2 maximum number of mismatches; −2 nonspecific match limit, and minimum fraction length of 0.7 for the genome mapping or 0.8 for the RNA mapping. The minimum reads similarity mapped on the reference genome was 80%. Only uniquely mapped reads were considered in the analysis. The libraries were normalized per million and the expression values for the transcripts were recorded in RPKM (Reads Per Kilobase per Million), we also analyzed the other expression values - TPM (transcripts per million) and CPM (counts per million). The statistical test applied was the DGE (Differential Gene Expression). For the ncRNA the database used was the ncRNA from *Histoplasma capsulatum* (EnsemblFungi G186AR GCA_000150115 assembly ASM15011v1). The secondary structure was performed using the PPFold plugin in the CLC Genomics Workbench v. 10.1 using the default parameters. Analysis of the relationship between the profile of RNA sequences detected in this study with the protein composition of *H. capsulatum* EVs was based on the results recently obtained with strain G217B using a proteomic approach (25). The cellular RNA used in this analysis was assessed from the SRA database (SRR2015219 and SRR2015223) (28).

### Data access

The data is deposited to the Sequence Read Archive (SRA) database under study accession number (PRJNA514312).

## Acknowledgments

JDN was supported in part by NIH R01AI052733 and R21AI124797. MLR is currently on leave from the position of Associate Professor at the Microbiology Institute of the Federal University of Rio de Janeiro, Brazil. He was supported by grants from the Brazilian agency Conselho Nacional de Desenvolvimento Científico e Tecnológico (CNPq, grants 405520/2018-2, 440015/2018-9 and 301304/2017-3) and Fiocruz (grants VPPCB-007-FIO-18 and VPPIS-001-FIO18). RP was supported by FAPESP (grant 13/25950-10). The authors also acknowledge support from Coordenação de Aperfeiçoamento de Pessoal de Nível Superior (CAPES, Finance Code 001) and the Instituto Nacional de Ciência e Tecnologia de Inovação em Doenças de Populações Negligenciadas (INCT-IDPN).

